# Colonic Epithelial miR-31 Associates with the Development of Crohn’s Phenotypes

**DOI:** 10.1101/307561

**Authors:** Benjamin P. Keith, Jasmine B. Barrow, Takahiko Toyonaga, Nevzat Kazgan, Michelle Hoffner O’Connor, Neil D. Shah, Matthew S. Schaner, Elisabeth A. Wolber, Omar K. Trad, Greg R. Gipson, Wendy A. Pitman, Matthew Kanke, Shruti J. Saxena, Nicole Chaumont, Timothy S. Sadiq, Mark J. Koruda, Paul A. Cotney, Nancy Allbritton, Dimitri G. Trembath, Francisco Sylvester, Terrence S. Furey, Praveen Sethupathy, Shehzad Z. Sheikh

**Author notes:** co-corresponding authors **Corresponding authors:** Terrence S. Furey PhD, Departments of Genetics and Biology, University of North Carolina at Chapel Hill, 5022 Genetic Medicine Building, 120 Mason Farm Road, Chapel Hill, NC 27599, Shehzad Z. Sheikh MD PhD, Department of Medicine and Genetics, University of North Carolina at Chapel Hill, 7340B Medical Biomolecular Research Building, 101 Mason Farm Road Chapel Hill, NC 27517, Praveen Sethupathy PhD, Department of Biomedical Sciences, College of Veterinary Medicine, Cornell University, T7010A 618 Tower Road, Ithaca, NY 14853.

## Abstract

Crohn’s disease (CD) is highly heterogeneous, due in large part to variability in cellular processes that underlie the natural history of CD, thereby confounding effective therapy. There is a critical need to advance understanding of the cellular mechanisms that drive CD heterogeneity. In this study, small RNA-sequencing and microRNA profiling in the colon revealed two distinct molecular subtypes, each with different clinical associations, in both adult and treatment-naïve pediatric CD patients. Notably, we found that microRNA-31 (miR-31) expression by itself can stratify patients into these two subtypes. Through detailed analysis of several colonic mucosa cell types from adult patients, we found that differential levels of miR-31 are particularly pronounced in epithelial cells. We generated patient crypt-derived epithelial colonoids and showed that miR-31 expression differences preserved in this *ex-vivo* system. In adult patients, low colonic miR-31 expression levels at the time of surgery are associated with post-operative recurrence of ileal disease. In pediatric patients, lower miR-31 expression at the time of diagnosis is associated with the future development of fibrostenotic ileal CD requiring surgery. These findings represent an important step forward in designing more effective clinical trials and developing personalized therapies for CD.

## Introduction

Crohn’s disease (CD), one of the primary inflammatory bowel diseases (IBD), is a chronic inflammatory condition of the gastrointestinal tract resulting from an aberrant immune response to the enteric microbiota in a genetically susceptible host. CD is highly heterogeneous in disease location, behavior, and progression. Using gene expression and chromatin accessibility profiles in colon tissue, we previously identified two molecular subtypes in adult CD associated with unique phenotypes (1). Recent studies validate the premise that specific genetic and molecular profiles are associated with, and may contribute to, disease heterogeneity and behavior. Over 200 genetic loci have been significantly associated with CD risk (2). A study of 29,838 adult individuals did not identify DNA variants predictive of CD behavior over time, but did associate genetic variants in IBD with disease location (3). Notably, a longitudinal inception cohort study of treatment-naïve pediatric CD patients revealed lipid metabolism and extracellular matrix gene expression signatures in the ileum as predictive of response to steroids and fibrostenotic ileal CD, respectively (4, 5). However, a more complete set of robust prognostic determinants for CD phenotypes, especially incorporating non-coding RNAs, is still lacking. As such, there remains active, substantive interest in the CD research community to identify specific genetic and molecular factors that mark disease subtypes, and more importantly, inform on disease progression and outcome.

Distinct disease outcomes of CD are likely due in large part to variability in cellular processes that underlie the natural history of CD. Disruption of the intestinal epithelial barrier and loss of tolerance by immune cells to the enteric microbiota are critical cellular events that lead to chronic inflammation seen in CD. Precise cell type-specific mechanisms leading to these dysfunctions are poorly understood. Recently, microRNAs (miRNAs) that confer post-transcriptional regulation of gene expression have emerged as key modulators of intestinal epithelial cell (IEC) biology (6, 7) and of pathways that underlie the pathogenesis of CD (8, 9). Mice deficient for miRNAs in the intestinal epithelium exhibit altered intestinal architecture and increased barrier permeability (6), which leads to immune cell infiltration and severe intestinal inflammation.

In this study, we identified miRNA-31 (miR-31) as the primary contributor to our previously identified two major molecular subtypes of adult CD patients. We determined that the upregulation of miR-31 in colonic tissue of CD patients is driven in large part by increased expression specifically in IECs. Importantly, we expanded our study to incorporate a large cohort of 234 formalin-fixed paraffin embedded (FFPE) index biopsies of colon and ileum tissue from 127 treatment-naïve pediatric patients and non-IBD (NIBD) controls. In medically refractory adult CD patients undergoing surgical resection, one subtype with a lower, more typical level of colonic miR-31 expression at the time of surgery was associated with a worse post-operative outcome (as measured by recurrence in the neo-terminal ileum at the anastomotic site) and need for subsequent colectomy. In pediatric patients, the same lower colonic miR-31 expression subtype in index biopsies was associated with progression to fibrostenotic ileal disease. Our study shows that miR-31 is a candidate prognostic determinant of CD behavior in adult and pediatric patients and highlights the potential role of miR-31 in the pathobiology of CD.

## Results

### MicroRNAs and lncRNAs stratify adult CD patients into two molecular subtypes

Previously, we demonstrated that medically refractory Crohn’s disease (CD) patients undergoing surgery clustered into two distinct groups using principal component analysis (PCA) of mRNA expression by RNA-seq on uninflamed colonic mucosa from 21 adult patients with CD and 11 adult control patients (NIBD) (1). Analysis of genes differentially expressed between these two groups revealed that genes more highly expressed in the colon of one group were enriched for previously identified NIBD colonic marker genes, while genes more highly expressed in the second group were enriched for normal ileum marker genes. We labeled these groups colon-like (CL) and ileum-like (IL). We showed by a prospective analysis that these CL and IL CD subgroups exhibit colonic CD and ileal inflammation, respectively. To evaluate further whether this molecular stratification was evident within non-coding RNAs, we analyzed small RNA-seq data from most of the same CD and NIBD patients (18 CD, 12 NIBD) to quantify the expression of microRNAs (miRNAs), which we previously showed was able to distinguish CD patients from NIBD controls (10). We also re-interrogated the RNA-seq data from the same patients to quantify long, non-coding RNAs (lncRNAs). PCA on each of the miRNA and lncRNA datasets (Figure 1A and 1B; Supplementary Tables 1 and 2) revealed that CD samples clustered into the same distinct CL and IL groups as initially defined with the mRNA data, which we also recapitulated in this study using updated gene annotations (Supplementary Figure 1 and Supplementary Table 3). These data demonstrate that the CL and IL CD subtypes are defined by expression profiles of several types of RNA molecules, which perform diverse functions within the cell.

**Figure 1.**
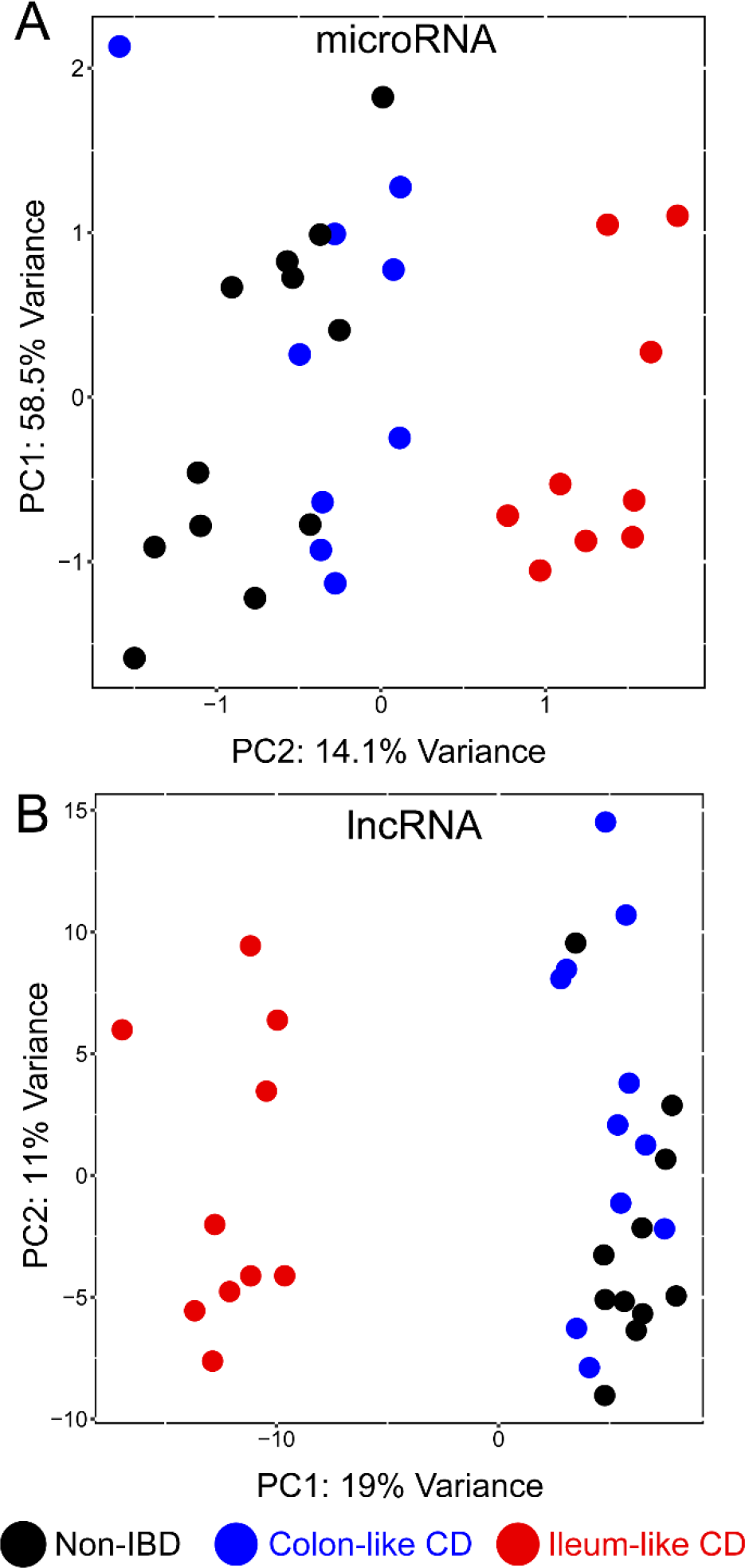
Two distinct molecular subtypes across multiple data types in adult Crohn’s disease (CD) Principal components analysis (PCA) of microRNA (A) and long non-coding RNA (B) expression profiles for patients with CD and patients with NIBD exhibit (black) distinct clusters; one enriched for colon-like CD patients (blue), and another enriched for ileum-like CD patients (red). Supplementary Tables 1 and 2 provide the PCA loadings for the above plots.

### miR-31 is the primary driver of molecular stratification and is associated with post-operative outcome in adult CD patients

To identify the miRNAs that contribute most to the stratification of the two molecular CD subtypes, we initially compared genome-wide miRNA expression profiles between the 9 CL and 9 IL CD patients. We found that 19 miRNAs were significantly differentially expressed between the two groups (|log_2_(FC)| > 1, FDR < 0.05). Strikingly, we observed a 13.5-fold change in miR-31-5p (miR-31; *P*_adj_ = 1.43 × 10^−18^) between CL and IL samples. Analysis of PCA components revealed that miR-31 is the top contributor to the variance observed for principle component (PC)-2 that separates the CL and IL patients (Supplementary Table 1). These findings suggest that miR-31 expression can stratify CD into two major molecular subtypes (Figure 2A).

**Figure 2.**
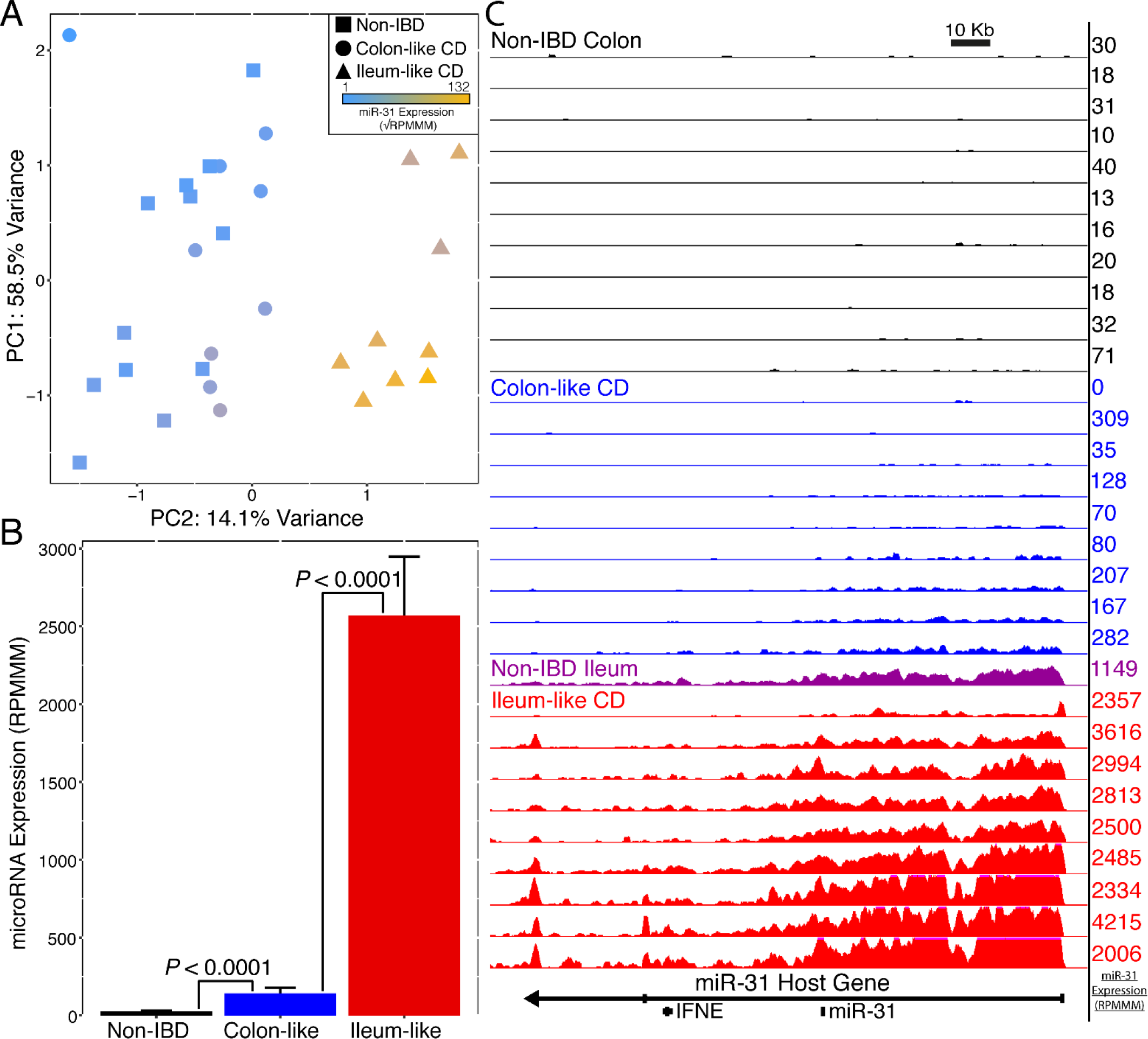
miR-31 is a driver of colon-like and ileum-like stratification. (A) Principal components analysis of miRNA expression data from small RNA-seq. miR-31 expression (blue-gold, low-high) appears to distinguish colon-like and ileum-like clusters. (B) Normalized miR-31 expression exhibits a significant upregulation in Crohn’s disease sub-groups samples compared with NIBD samples. (C) UCSC browser representation of normalized RNA-seq reads mapping to the miR-31 host gene for NIBD colon (black), colon-like CD (blue), NIBD Ileum (purple) and ileum-like CD (red). miR-31 transcript expression levels from small RNA-seq (RPMMM) are displayed to the right of each track. FDR adjusted p-values determined using DESeq2, with data presented as mean RPMMM ± SE.

We and others have identified miR-31 as a discriminant more generally of CD and NIBD patients (11, 12). We hypothesized that this difference is driven primarily by CD patients in the IL group. To test this hypothesis, we compared the levels of miR-31 in each of IL and CL groups relative to NIBD. We observed a dramatic and highly significant up-regulation (~60-fold) of miR-31 in IL patients compared with NIBD patients (*P*_adj_ = 2.59 × 10^−51^; Figure 2B). We also detected a significant difference in expression between CL and NIBD (~4-fold, *P*_adj_ = 7.66 × 10^−06^; Figure 2B); however, the magnitude of the difference is much lower. These findings support the above-stated hypothesis, indicating that while miR-31 is a strong marker of disease presence in all CD patients, this signal is driven predominantly by those patients of the IL subtype.

Expression levels of mature miRNAs can be altered in several ways, including changes to the rate of transcription, efficacy of the maturation (biogenesis) process, and RNA stability. To determine whether the miR-31 locus is subject to enhanced transcription in IL CD patients, we quantified the normalized density of RNA-seq reads mapping to the primary transcript of miR-31 (MIR31HG) across all samples. We observed that transcription levels of MIR31HG are indeed dramatically elevated in the IL subgroup relative to both the CL subgroup and NIBD patients (Figure 2C). These data indicate that increased level of transcription is one major contributor to the observed difference in miR-31 levels between IL patients and the NIBD and CL patients. Notably, RNA-seq data from the ileum of an NIBD patient (Figure 2C) revealed a signal at the MIR31HG locus that closely resembles the signal from the colon of IL CD patients.

MiRNAs regulate gene expression by binding to recognition elements in the 3’ untranslated regions of target mRNAs and marking the mRNAs for translational repression and degradation (13). Therefore, we sought to determine, using our published tool miRhub (14), whether genes that are down-regulated in IL relative to NIBD are enriched for predicted target sites of miR-31 or any other miRNA shown to be upregulated in the colon of IL patients. Notably, we found that miR-31 is the only upregulated miRNA whose target genes are significantly enriched among the genes downregulated in IL patients compared to both CL and NIBD patients (empirical *P* < 0.05; Supplementary Figure 2). This indicates that miR-31 is not only dramatically elevated in the IL subtype of CD, but also a candidate master regulator of genes that are downregulated in that subtype.

To validate the differential expression of miR-31 between the IL and CL subtypes of CD, we measured colon miR-31 levels in an independent cohort of 67 adult CD and 42 NIBD patients using qRT-PCR. Biopsies were obtained at the time of surgical resection for medically refractory disease. We first recapitulated the finding that miR-31 levels are significantly up-regulated overall in CD relative to NIBD (*P* = 1.72 × 10^−4^, 2-tailed unpaired Student’s *t* test; Figure 3A). As expected, we also found that miR-31 expression levels stratify CD patients into two subgroups, “high” and “low”, which we hypothesized reflect the IL and CL molecular subtypes, respectively. To test this hypothesis, we measured mRNA levels of *APOA1* (Fig 3B), a marker gene in ileum, and *CEACAM7* (Figure 3C), a marker gene in colon, both of which we previously showed can stratify IL and CL patients (15, 16). We found that the patients with high colonic miR-31 expression also show high *APOA1* expression and low *CEACAM7* expression, and we observed the opposite trend for the patients with low colonic miR-31 expression. Although baseline demographic and clinical characteristics of CL and IL subtype patients at the time of miR-31 measurement were not statistically different, there was a trend towards the increased presence of ileal stricturing in the IL subtype (Table 1). Altogether, these data confirm that miR-31 expression levels stratify CD patients into two molecular subgroups.

**Figure 3.**
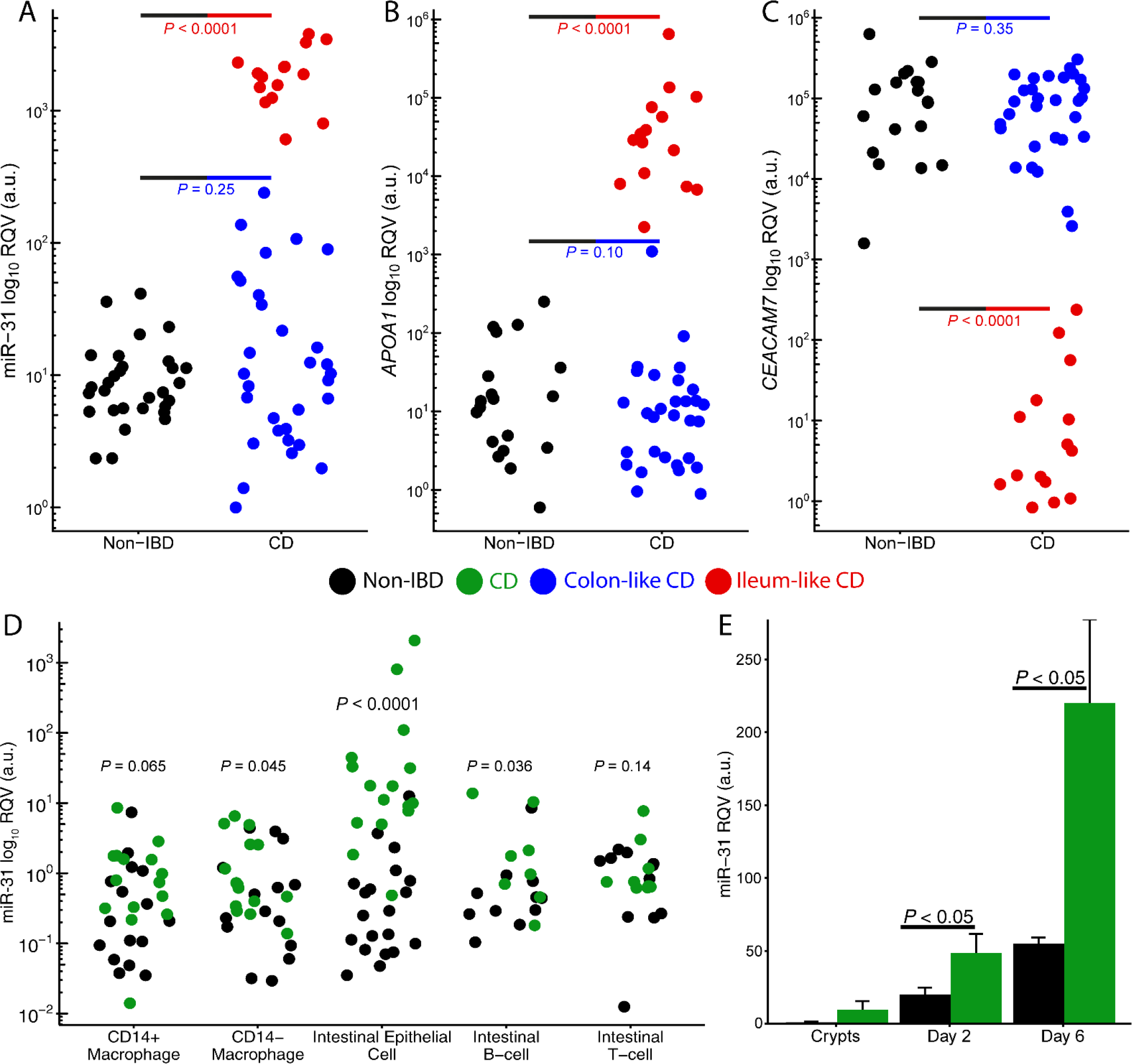
miR-31 is specifically upregulated in intestinal epithelial cells. qRT-PCR for miR-31 (A), APOA1 (B), and CEACAM7 (B) in an independent adult cohort displays colon-like (blue; n=42) and ileum-like (red; n=25) clustering patterns for CD samples compared with NIBD samples (black; n=42). (D) qRT-PCR of five colon-specific cell types reveal significant miR-31 upregulation in intestinal epithelial cells isolated from CD patients (n=11-20) relative to NIBD controls (n=8-16). (8 NIBD matched and 6 CD matched across all cell types). (E) Relative miR-31 expression by qPCR of in colonoid cultures generated from NIBD controls (n=4) compared with CD patients (n=4). miR-31 expression is increased in fresh crypts and remains higher at day 2 and day 6 of colonoid culture. miRNA levels are relative to RNU-48 expression compared to fresh NIBD crypts. Significance values determined by a 2-tailed unpaired Student’s t test. Data are presented as mean ± SE.

**Table 1.** Demographics of adult CD patients. CD patients were classified into colon-like and ileum-like CD molecular subtypes. Phenotype associations with subtypes were assessed using Fisher’s exact test (categorical data) and 2-tailed unpaired Student’s t test (continuous data). *P* < 0.05 (bolded) was considered statistically significant. Individual patient demographics are presented in Supplementary Table 4.

| Phenotype | Colon-like<br>(n=42) | Ileum-like<br>(n=25) | P Value |
| --- | --- | --- | --- |
| **Patient Characteristics** |  |  |  |
| Age at Sample Collection (years) | 37.95 | 34.28 | 0.327 |
| Male | 20 | 12 | 1.000 |
| Female | 22 | 13 | 1.000 |
| Smoker (current or previous) | 16 | 9 | 1.000 |
| **Location** |  |  |  |
| Ileum-only | 9 | 4 | 0.753 |
| Colon-only | 11 | 3 | 0.222 |
| Ileum+Colon | 22 | 18 | 0.131 |
| Upper GI | 7 | 4 | 1.000 |
| **Phenotypes and Involvement** |  |  |  |
| Perianal | 18 | 6 | 0.187 |
| Ileal Disease | 31 | 22 | 0.222 |
| Inflammatory | 7 | 0 | **0.040** |
| Strictureing | 19 | 18 | **0.044** |
| Penetrating | 5 | 4 | 0.072 |
| Disease Duration (years) | 12.43 | 10.64 | 0.517 |
| **Pre-operative treatment history** |  |  |  |
| Steroids | 25 | 13 | 0.615 |
| 5-ASA | 12 | 11 | 0.287 |
| Immunomodulation | 12 | 13 | 0.070 |
| Anti-TNF | 23 | 15 | 0.800 |
| Non-anti-TNF biologic | 6 | 1 | 0.244 |

We then studied prospectively the clinical characteristics of adult patients after surgery for medically refractory disease. Since all patients had disease removed at initial surgery, we defined recurrence of disease based on Rutgeerts post-operative endoscopic scoring (17) of the neo-terminal ileum or the need for an end ileostomy due to severe refractory disease within a year after the initial surgery. Post-operative management as well as timing of endoscopy for reassessment was determined by the managing IBD specialist. Most patients (Supplemental Table 4) had a post-operative staging colonoscopy within one year of surgery. Recurrence was defined as having a Rutgeerts score of i2, i3, i4 or the need for an end ileostomy within a year after the initial surgery. No recurrence was defined as a Rutgeerts score if i0, i1. Strikingly, despite similar patient demographics at time of surgery as well as no significant differences in post-operative management between the two subtypes (Table 2), the CL subtype of CD patients demonstrated a worse post-operative course compared to the IL subtype (Table 2, p=0.024). While, this patient population is not anti-TNF treatment naïve, to our knowledge, this data provides the first evidence for the potential clinical utility of miRNA profiling to predict a poor post-operative outcome of CD.

**Table 2.** Post-operative clinical characteristics of adult CD patients. Associations between CD molecular subtypes with post-operative phenotypes were assessed using Fisher’s exact test (categorical data) and 2-tailed unpaired Student’s t test (continuous data). *P* < 0.05 (bolded) was considered a statistically significant association. Recurrence was defined as having a Rutgeerts score of i2, i3, i4 or the need for an end ileostomy within a year after the initial surgery. Remission was defined as a Rutgeerts score if i0, i1. Adult phenotypes for individual patients are shown in Supplementary Table 4.

| Phenotype | Colon-like<br>(n=42) | Ileum-like<br>(n=25) | *P* Value |
| --- | --- | --- | --- |
| Surgery | 36 | 25 | 0.077 |
| Post-operative Anti-TNF Use | =23/36 | =16/25 | 1.000 |
| End ileostomy | =11/36 | =1/25 | **0.019** |
| Second resection | =17/36 | =10/25 | 0.610 |
| Time to first resection (years) | 6.86 | 8.20 | 0.481 |
| Time from first to second resection (years) | 11.00 (=16/36) | 5.70 (=10/25) | 0.25 |
| Post-op Colonoscopy | =35/36 | =20/25 | NA |
| Remission | =11/35 | =13/20 | **0.024** |
| Recurrence | =24/35 | =7/20 | **0.024** |

### miR-31 is dramatically up-regulated in intestinal epithelial cells and crypt derived colonoids established from adult CD patients

Colon tissue is composed of several distinct cell types, and expression studies in tissue do not reveal from which particular cells transcripts originated. To measure miR-31 expression in specialized cell types of the colon, we isolated intestinal epithelial cells (IECs; CD326+) and matched lamina propria immune cells (CD3+ T cells, CD20+ B cells, CD33+CD14-resident intestinal macrophages, CD33+CD14+ infiltrating inflammatory intestinal macrophages) by flow cytometry (Supplementary Figure 3) from macroscopically uninflamed tissue from adult patients with CD (N=11-20) and NIBD controls (N=8-16). While relative miR-31 expression levels based on qRT-PCR were increased in B cells and resident macrophages isolated from CD patients compared to NIBD controls (P < 0.05, 2-tailed unpaired Student’s t test), these results were dwarfed in comparison to the increase seen in IECs (~52-fold difference, P = 1.28 × 10^−8^; Figure 3D).

To evaluate this finding further, we established three-dimensional epithelial colonoids from crypts isolated from both CD patients and NIBD individuals. These structures contain crypt-like domains reminiscent of the gut epithelium, and they continuously produce all cell types found normally within the intestinal epithelium (18). We found that colonoids from CD patients express significantly higher levels of miR-31 compared to NIBD controls, similar to the primary tissue from which the colonoids were derived (Day 2 *P* = 0.041, 2-tailed unpaired Student’s t test; Day 6 *P* = 0.0095, 2-tailed unpaired Student’s t test; Figure 3E). These results suggest upregulated miR-31-5p is not a transient result due to external signalling but is a predisposing factor in IECs of CD patients. Disruption of the intestinal epithelial barrier is a critical determinant of the predisposition to chronic inflammation and fibrosis seen in CD. Going forward these data open up the potential to understand the impact of miR-31 on barrier function.

### miR-31 expression in formalin-fixed paraffin-embedded (FFPE) tissue from treatment-naïve pediatric CD patients also defines two subtypes and is associated with development of ileal fibrostenotic disease

The molecular profiles we have generated and analyzed in fresh tissue and cells from adult CD represent a fundamental advance in understanding adult CD heterogeneity. At the time of this analysis, though, these adult patients had progressed to medically refractory disease, each with individual treatment histories that could potentially confound results. Therefore, as a next step, we performed smRNA-seq on microscopically uninflamed FFPE mucosal tissue from ascending colon and terminal ileal biopsies in age-matched treatment-naïve pediatric patients with CD (n=76) and NIBD controls (n=51) obtained at the time of diagnosis (index colonoscopies). It is important to note that this is not a validation cohort of the adult CD, but rather a completely independent analysis that offers at least five unique advantages. Firstly, as noted above, these samples are from treatment-naïve individuals, which greatly mitigates the potential confounding effects of treatment history that may be present in adults. Secondly, the samples are FFPE as opposed to fresh frozen tissue. Successful molecular subtyping of CD patients using FFPE tissue will greatly expand our ability in the future to analyze retrospectively the clinical characteristics associated with subtypes, given that most tissue biopsies are bioarchived as FFPE. Thirdly, the number of samples is substantially greater than in our adult CD study, affording additional power for molecular subtyping. Fourthly, we have matched ileum and colon biopsies from the same patient allowing for the interrogation of site specific changes and impact on disease phenotype. Finally, these tissue samples are index biopsies, obtained at the time of diagnosis and prior to significant disease progression, which provides a unique opportunity to determine whether miR-31 expression is associated with the development of CD phenotypes.

As in the adult cohort, we found that the levels of miR-31 expression in the colon are significantly upregulated in CD patients relative to NIBD controls (~7.8-fold, *P* = 4.64 ×10^−7^, 2-tailed unpaired Student’s *t* test; Figure 4A and Supplementary Figure 4). We observed that miR-31 expression in the ileum is also significantly upregulated in CD patients (*P* = 9.97 × 10^−7^, ~1.5-fold), however the effect is not nearly as pronounced as in the colon (Figure 4B). This may be due in part to significantly higher baseline miR-31 expression levels in the ileum of unaffected (NIBD) individuals compared to in the colon (*P* = 5.71 × 10^−28^; Supplementary Figure 5 and Supplementary Table 5).

**Figure 4.**
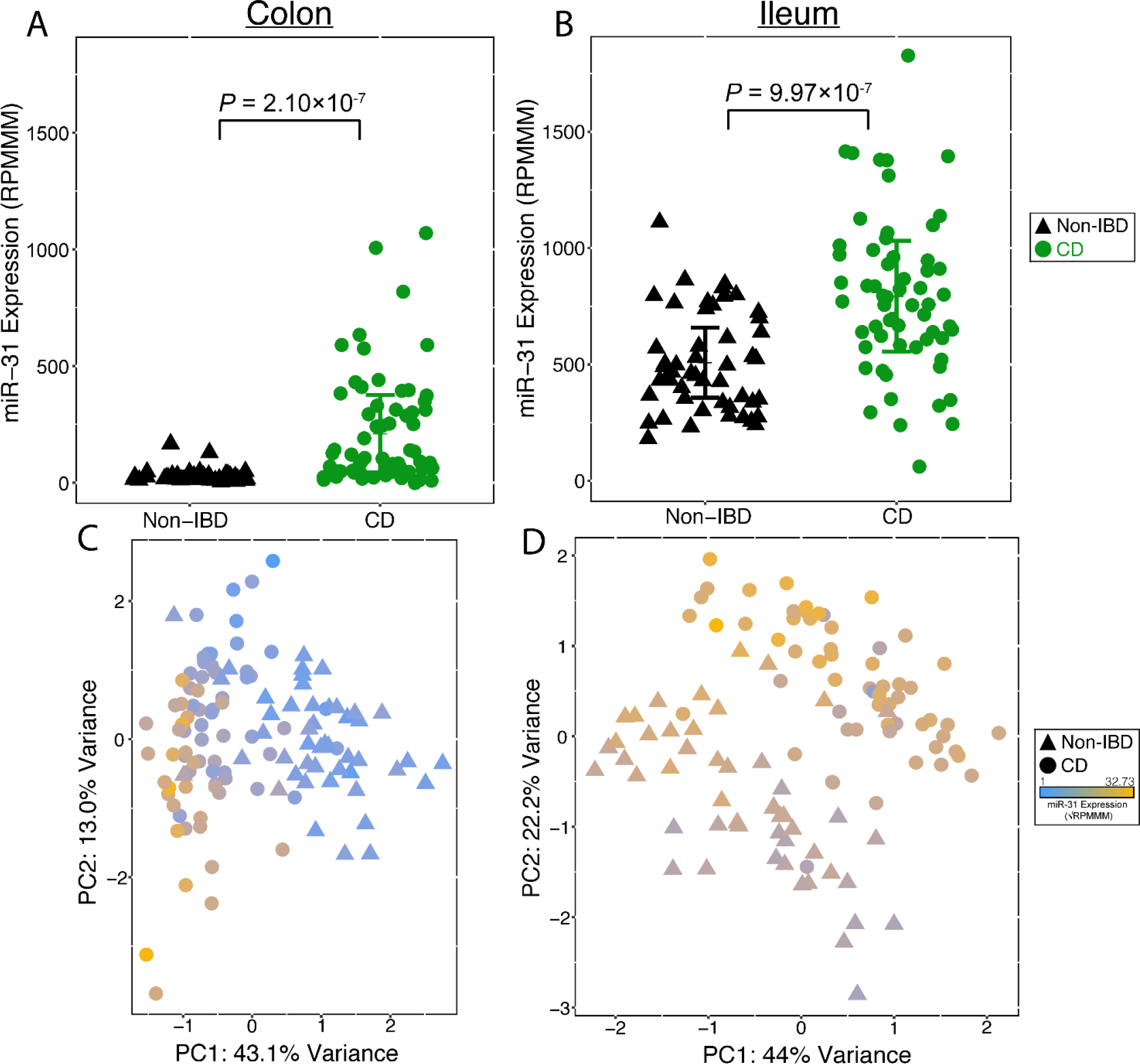
miR-31 is differentially expressed in treatment-naïve pediatric Crohn’s disease (CD) samples. miR-31 expression is significantly upregulated in the colon (A) and ileum (B) of treatment-naïve pediatric samples compared with pediatric NIBD samples. Principal components analysis of miRNA expression profiles from small RNA-seq results in distinct clusters of NIBD and CD patients for colon (C) and ileum (D) samples. Points are colored according to miR-31 expression (blue-gold; low-high). Significance determined by a 2-tailed unpaired Student’s t test where P < 0.05. Data presented as mean ± SE

Using miRNA expression data from the 100 most variable miRNAs, we independently performed PCA on the colon (Figure 4C) and ileum (Figure 4D) pediatric samples and observed a robust separation of NIBD and CD patients. Notably, miR-31 is the largest contributor to this stratification in the colon, but not in the ileum (Supplementary Tables 6 and 7). This indicates that specifically colonic miR-31 is a primary marker of disease presence.

We investigated whether colonic miR-31 levels were associated with the eventual development of specific CD phenotypes and tested for association with clinical features both at the time of diagnosis and across disease course (Table 3). We first analyzed pediatric NIBD samples and found that all colon samples but one had miR-31 levels < 150 RPMMM and all ileum samples had mir-31 levels > 150 RPMMM (Supplementary Figure 6). Using this threshold, we defined two distinct subgroups within our colonic pediatric CD samples as “miR-31-low” (n = 46) and “miR-31-high” (n = 30). MiR-31 expression was validated in our two subgroups through qRT-PCR of a subset of low- (n = 7) and high-miR-31 (n = 7) samples (*r* = 0.94, *P* = 3.89 × 10^−7^; Supplementary Figure 7).

**Table 3.**
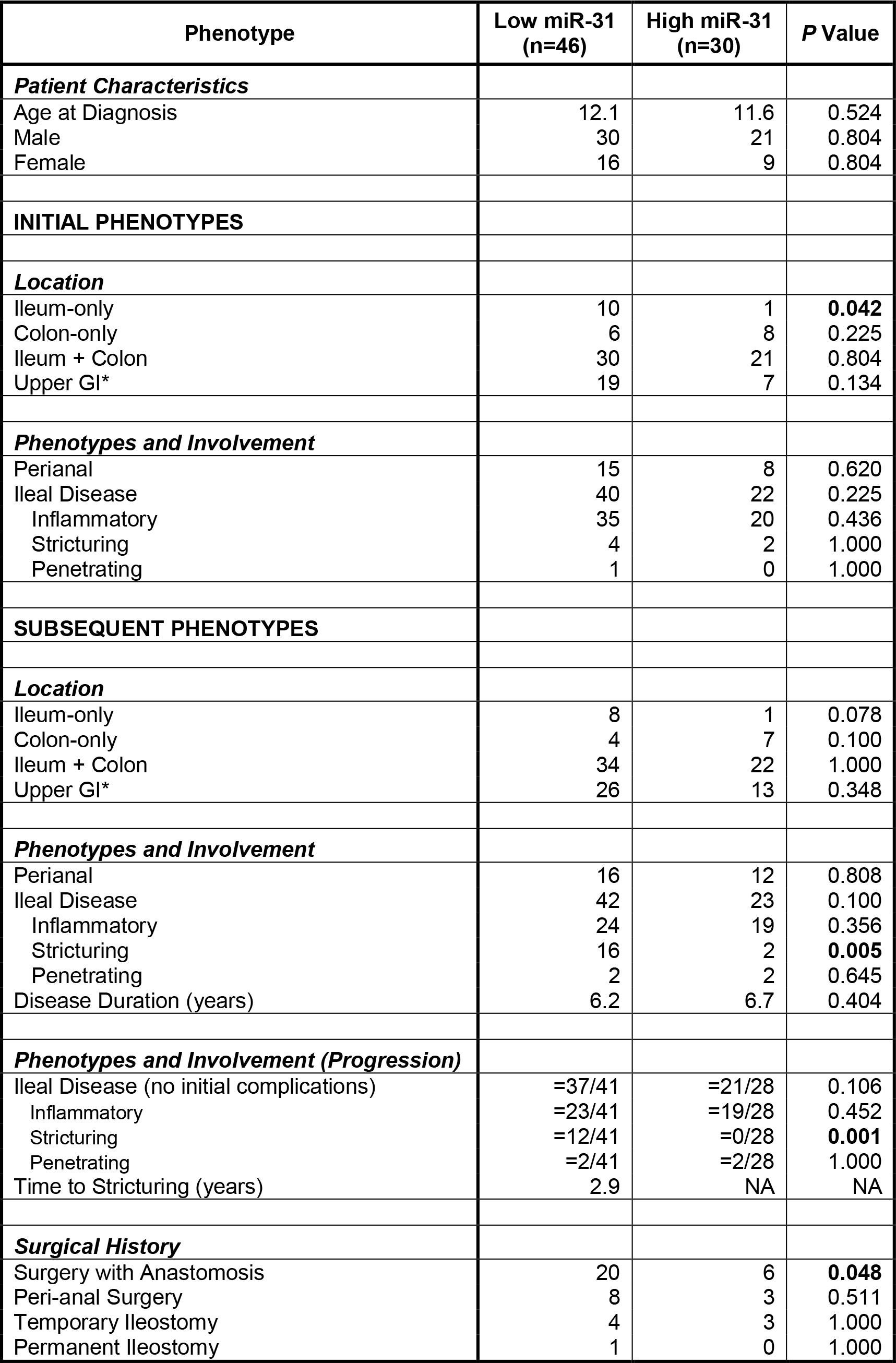
Clinical phenotypes of pediatric CD patients. Pediatric CD patients were classified into low miR-31 expression (<150 RPMMM; n=46) and high miR-31 expression (≥150 RPMMM; n=30) groups. Clinical phenotypes were recorded at time of initial diagnosis when miR-31 expression was determined, and at subsequent time points after these initial diagnoses. Location of disease in the upper gastrointestinal (GI) tract is in addition to colonic and/or ileal disease. Only patients that initially presented with inflammation only and no complications were considered when assessing progression to disease complications. Associations between molecular subtypes and clinical phenotypes were assessed using Fisher’s exact test and were performed only on categories with at least 8 patients across both subtypes. Significant associations (p<0.05) are bolded. Pediatric phenotypes for individual patients are shown in Supplementary Table 8.

We then studied prospectively the clinical characteristics of only pediatric patients that presented with inflammation at time of diagnosis (i.e., no initial stricturing, penetrating disease). Since all patients were treatment naïve, we defined stricturing CD as primary (not anastomotic) fibrostenotic stricture of the terminal ileum where medical treatment would be ineffective, and therefore, surgical resection was considered a reasonable treatment option. These were diagnosed based on physician preference of using standard endoscopy and/or computed tomography (enterography) (CTE) or magnetic resonance imaging (MRI) and correlation with patient symptoms. Low miR-31 expression was significantly associated with the eventual development of ileal stricturing (*P* = 0.001) and having surgery involving an anastomosis (*P* = 0.048). Remarkably, we found that no miR-31-high patients progressed to develop a stricturing phenotyping. To our knowledge, this data provides the first evidence for the potential clinical utility of miRNA profiling to predict increased risk of the development of stricturing phenotype in patients with Crohn’s disease.

## Discussion

We identified colonic miR-31 expression as central to clinically-relevant molecular subtypes found in independent cohorts of adult and treatment-naïve pediatric patients. Notably, low levels of miR-31 in medically refractory adult CD patients at the time of surgical resection are indicative of a worse post-operative outcome as measured by recurrence in the neo-terminal ileum. Similarly, lower miR-31 expression in pediatric patients at the time of diagnosis is indicative of increased risk for development of ileal stricturing complications. Our study introduces small RNAs as potential predictors of disease phenotype and, with use of FFPE samples, offers distinct advantages over mRNA studies in the context of fresh tissue. These findings are reminiscent of early descriptions of transcriptomic signatures in breast cancer (19). Further large-scale studies of gene expression profiles in breast tumors, including those of The Cancer Genome Atlas (TCGA) project (20), eventually established four major molecular classes that vary in their aggressiveness and respond differently to therapies. Similarly, diffuse large B cell lymphoma (21), glioblastoma (22), endometrial cancer (23), and lung cancer subtypes (24) have been identified by genomic profiling, facilitating the development and application of targeted therapies (https://cancergenome.nih.gov).

Our study includes two distinct populations of patients with disease at different stages of development. It is unclear how clinical associations are related to patient age and/or disease state. For instance, molecular levels at initial diagnosis that predict disease progression may not be maintained once disease has actually progressed (25). Long-term longitudinal studies will need to be conducted with serial quantification of genomic profiles over the course of disease evolution in pediatric patients transitioning into adulthood. It is also imperative we understand the unique characteristics of disease presentation and evolution in adult patients, impacted by major life style and environmental factors, each uniquely contributing to colonic miR-31 regulation and its impact on phenotype. Medical management of fibrostenotic ileal disease is unpredictable and in many cases, is not long lasting and requiring surgery. Thus, results from these longitudinal studies may eventually impact treatment designs for these difficult disease phenotypes. The robust establishment of CD subtypes may also influence future design of clinical trials where subtypes can be considered during patient randomization, allowing for better evaluation of subtype identification when making therapeutic decisions.

Recent studies have started to unravel the molecular mechanisms associated with distinct IBD phenotypes. Genetic variants in NOD2, MHC, and MST1 3p21 were shown to be associated with disease location (colonic CD, ileal CD and ulcerative colitis (UC)) but not disease behavior (3). But, the genetic contribution to CD pathogenesis has been shown to be disproportionate, ranging from most impactful in very early onset IBD (26) (VEOIBD) to modest significance in older pediatric and adult IBD patients (27–30). In rectal tissue from pediatric patients, expression patterns of IL-13, IL23A, and IL17 distinguished colonic CD from UC (31). Also, a lipid metabolism related gene expression signature in the ileum of pediatric CD patients accurately predicted 6-month steroid-free remission (4). Follow-up studies of these same ileal samples showed a distinct collagen and extracellular matrix gene expression signature present at time of diagnosis in a subset of patients who developed fibrostenotic ileal disease (5). Interestingly, our prior analysis of these patients identified an association between these same pathways in the ileum and the CL molecular phenotype (1). Moving forward, the challenge is to define molecular subtypes while also uncovering the cell type-specific genetic, molecular, and environmental contributors to each subtype.

This current study along with our previous study have now shown that whole genome mRNA, miRNA, and lncRNA transcript levels, along with the open chromatin landscape, define two molecular adult CD subtypes. In addition, miRNA expression patterns can stratify pediatric CD. Together, these findings suggest that across CD patients, colonic tissue is altered in different ways at a cellular level supporting the idea of multiple Crohn’s diseases. This also underscores the necessity for a more complete molecular characterization of CD across larger populations to uncover additional distinct subtypes. We advanced our work into FFPE tissue which opens the possibility to increase sample numbers, perform longitudinal follow-up studies, and facilitate the association of molecular markers to disease course.

We demonstrate miR-31 to be specifically dysregulated in colonic epithelial cells. Breakdown in the intestinal barrier is critical to intestinal chronic inflammation; a hallmark of CD. MiRNAs, including miR-31, are known to have significant contributions to gastrointestinal epithelial barrier function (32). *Dicer1* deficient mice display colonic barrier integrity dysfunction as evidenced by lymphocyte and neutrophil infiltration as well as mis-localization of the tight junction protein Claudin-7 (6). In the esophagus, Hussey et al. found that miR-31 is one of only a few differentially expressed miRNAs in post-ablation epithelium with increased barrier permeability (33, 34). In the colon, Wu et al. postulate that lowly expressed miR-31 plays a protective role after hypothermic ischemia induced barrier dysfunction in the colon, perhaps aiding in post-injury healing, specifically by targeting the hypoxia inducible factor (HIF)-factor inhibiting HIF (FIH-1) pathway (35). Using combinational computational methods to predict miR-31 target-pathways, one group found a connection specifically between miR-31 and tight junctions in lung epithelium (36). Most recently, Yu et al. demonstrated using *in vivo* knock-in and knock-out models that miR-31 plays a role in regulating intestinal stem cell behavior during regeneration after radiation injury (37). We show that patient crypt-derived colonoids in a sterile environment retain the aberration in miR-31 expression present in the tissue of origin, which supports a cellular defect that is intrinsic and not secondary to inflammation or other external signals due to the presence of disease. The colonoid experimental system will enable future studies to interrogate the role(s) of specific factors in driving a fibrostenotic phenotype, especially in the context of co-culture with lamina propria immune cells, mesenchymal cells, as well as stimulation with commensal and/or colitogenic bacteria.

In summary, we provide the most comprehensive molecular characterization of CD to date. We uncover miR-31 as an identifier of CD, but more importantly a molecular stratifier of both pediatric and adult patients, an indicator of established disease phenotype in adult patients, and a predictor of clinical phenotype at the time of diagnosis in pediatric patients. These findings represent significant progress in molecularly defining the Crohn’s disease(s), moving closer toward potential personalization of therapy and improving outcomes.

## Methods

### Patient populations and phenotyping

Adult and pediatric patients with CD and NIBD related illnesses diagnosed at The University of North Carolina hospitals (UNC) were included in this study. Clinical phenotypes considered in this study include demographic and clinical variables such as age, sex, disease duration, age at diagnosis, age at sample acquisition, disease location, and disease behavior. Summarized (Supplementary Table 9) and detailed information of patient demographics and phenotypes for the adult (Supplementary Table 4) and pediatric (Supplementary Table 8) cohorts are provided. This study was not blinded, and all authors had access to the study data and reviewed and approved the final manuscript.

### Tissue isolation and characterization

For our adult cohort, all CD and NIBD mucosal biopsies were obtained from macroscopically unaffected sections of the ascending colon at the time of surgery and flash-frozen. No samples showed signs of active microscopic inflammation or disease, as confirmed by an independent pathologist (DGT). Treatment-naïve pediatric patients were diagnosed at UNC. From formalin-fixed, paraffin-embedded (FFPE) tissue, mucosal sections from both macroscopically and microscopically non-inflamed sections of the ascending colon and terminal ileum from the time of initial diagnosis (index biopsy) were identified by a pathologist (DGT), and scrolls were obtained for small RNA isolation. Absence of acute (active) inflammation, including neutrophilic inflammation of crypt epithelium and crypt abscess formation, and chronic inflammation, including architectural distortion and basal lymphoplasmacytosis of the lamina propria, was determined after review of each H&E stained slide (Supplementary Figure 8).

### RNA isolation, sequencing, and analysis

RNA was isolated from flash-frozen adult samples from surgical resections using the Qiagen RNeasy Mini Kit (Valencia, CA) following the manufacturer’s protocol. This kit uses column-based DNase treatment to eliminate DNA contamination, and allows the miRNA and mRNA content to be preserved. miRNA was enriched from FFPE tissue for pediatric samples using the Roche High Pure miRNA Isolation Kit (Penzberg, Germany). RNA purity and integrity were assessed with Thermo Scientific NanoDrop 2000 (Waltham, MA) and Agilent 2100 Bioanalyzer (Santa Clara, CA), respectively. For all clinical categories of flash frozen adult samples, we observed average RNA integrity (RIN) values above 7.

RNA-seq libraries were prepared using the Illumina TruSeq polyA+ Sample Prep Kit. Paired-end (50 bp) sequencing was performed on the Illumina HiSeq 2500 platform (GEO accession GSE85499). Reads were aligned to the GRCh38 genome assembly using STAR (38) with default parameters. Transcript expression was quantified with Salmon (39) using default parameters. Post-alignment normalization and differential analysis was performed using DESeq2 (40) with GENCODE_V25 gene annotations requiring base mean expression >10 and an FDR <0.05.

Small RNA libraries were generated using Illumina TruSeq Small RNA Sample Preparation Kit (San Diego, CA). Single-end (50 bp) sequencing was performed on the Illumina HiSeq 2500 platform (GEO accession GSE101819). miRquant 2.0 (41) was used for miRNA annotation and quantification. Samples with less than 3 million reads mapping to miRNAs were excluded. Differential analysis was performed using DESeq2 (40).

PCA was performed using the prcomp function in R on DESeq2 normalized VST transformed counts for mRNAs (“protein_coding” in GENCODE_V25) and lncRNAs (“lincRNA” or “antisense” in GENCODE_V25) with an expression base mean > 10. For miRNA expression data, PCA was performed using reads per million miRNAs mapped (RPMMM) normalized log2 transformed counts for the 100 miRNAs with the highest standard deviation values across all samples and a normalized expression level of 500 RPMMM across at least 20% of samples. For pediatric samples, we eliminated 18 miRNAs not found in the adult samples to remove potential artifacts due to FFPE preservation. Candidate master regulator miRNAs were detected using miRHub (14), using “non-network” mode and requiring a predicted target site to be conserved between human and at least two other species.

### Quantitative reverse transcriptase PCR

For miR-31, total RNA was isolated from tissues using Norgen’s Total RNA Purification Kit (Thorold, ON, Canada). 50ng of RNA was used for reverse transcription with the Life Technologies TaqMan MicroRNA Reverse Transcription Kit (Grand Island, NY). MiRNA qRT-PCR were performed using the TaqMan Universal PCR Master Mix per Life Technologies’ protocol, on Bio-Rad Laboratories CFX96 Touch Real Time PCR Detection System (Richmond, CA). Reactions were performed in triplicate using RNU48 as the normalizer. For APOA1 and CEACAM7, total RNA was isolated as described above. cDNA was derived from 1μg RNA by reverse transcriptase using the BioRad iScript cDNA Synthesis kit. RT-qPCR was then performed on these cDNA samples using the BioLine Hi-ROX SYBR kit.

LPMCs and IECs were isolated from intestinal specimens using modifications of previously described techniques (42). LPMCs were isolated from human colon by an enzymatic method, followed by Percoll (GE Healthcare, Piscataway, NJ) density-gradient centrifugation. LPMCs were further separated into CD33+14+ peripheral macrophages, CD33+CD14-intestinal resident macrophage, CD20+ B cells, and CD3+ T cells corresponding antibody labeled microbeads (Miltenyi Biotec, Auburn, CA). IECs were isolated from human colon mucosa using Ethylenediaminetetraacetic acid (EDTA) followed by magnetic bead sorting via CD326 labeled microbeads. Purity was >90% by flow cytometric analysis (Supplementary Figure 3)

### Colonoid generation and analysis

Epithelial colonoid cultures were generated from non-inflamed regions of colon tissue from NIBD controls and CD patients. The intestinal tissues were washed and mucosectomy performed with surgical scissors. Minced colonic mucosal fragments were incubated at 37°C in 5 ml of digestion media (1 mg/ml collagenase VIII in Advanced Dulbecco’s modified Eagle medium/F12 (ADF), 10% FBS, 15mM HEPES buffer, penicillin/streptomycin, 2mM Glutamax, 100ug/ml Primocin (Invivogen, antibiotic/antimitotic), 10uM Y-27632) for 30 minutes with mechanical disruption. The digested tissue/crypts were centrifuged at 200*g* for 5 minutes to separate crypts from single cells. Pelleted colonic crypts were resuspended in 5 ml of digestion media and centrifuged again at 200*g* for 5 minutes. Volume of crypts needed for 40-50 crypts per 96-well well was centrifuged in 1.5 mL tubes at 2500 RPM for 5 minutes. Crypts were embedded in appropriate volume of Growth Factor Reduced Matrigel (Corning) on ice and seeded at 10uL per 96-well. Basal stem culture medium (50% WNT3a conditioned media, 50% R-spondin 2 conditioned media, supplemented with 1 mM HEPES, 2mM Glutamax, 1X N2, 1X B27, and 1 mM *N*-acetylcysteine, 100ug/ml Primocin, with growth factors 50ng/mL murine EGF, 100ng/mL murine noggin, 1 ug/mL gastrin, 0.01uM PGE2, 10mM nicotinamide, and small molecule inhibitors 500 nM LY2157299, 10 uM SB202190) with 10 uM Y-27632 was added at 100uL per well. At selected timepoints, colonoids embedded in matrigel were lifted from wells with cold ADF. For miRNA analysis, day 2 and 6 reverse transcription and quantitative real time PCR for miR-31 and RNU-48 (housekeeping) were performed using predesigned TaqMan miRNA assays (Life Technologies). The relative expression was calculated by the comparative CT method and normalized to the expression of RNU-48.

### Statistics

Differential expression analyses of RNA-seq and small RNA-seq data were performed using DESeq2 (40), with FDR adjusted p-values being used to measure statistical significance. MicroRNA target enrichment was determined using miRHub, which generates empirical p-values through Monte Carlo simulations (14). Significance of differential expression in RT-qPCR and colonoid assays was assessed using Student’s t test (unpaired, 2 tailed) to compare 2 groups of independent samples. Significance of association with patient phenotype data was determined using Fisher’s exact test (categorical data) or a 2-tailed unpaired Student’s t test (continuous data). For all tests, *P*_adj_ < 0.05, empirical *P* < 0.05, or *P* < 0.05 was considered statistically significant.

### Study Approval

Both the adult and pediatric sections of this study received Institutional Review Board approval at UNC (protocol 10-0355 and 15-0024). Written informed consent was received from all participants prior to inclusion in the study. All participants are identified by number and not by name or any Protected Health Information (PHI).

## Author Contributions

BPK acquired, analyzed and interpreted data, prepared figures, drafted and revised the manuscript. JBB acquired, analyzed and interpreted data and revised the manuscript. NK acquired, analyzed and interpreted data; GRG, MSS, MH, SSS, OKT, PAC, TT, and NA acquired data; and WAP and MK analyzed data. NDS, EAB, NS, TSS, MJK, DGT and FS provided help with tissue acquisition and patient phenotyping. TSF & PS designed the study, analyzed and interpreted the data, drafted and revised the manuscript, and obtained funding. SZS conceptualized and designed the study, acquired the data, interpreted data, drafted and revised the manuscript, obtained funding, acted as study sponsor, and supervised the study. All authors uphold the integrity of the work, have had final approval of the manuscript in its entirety, and are accountable for all aspects of the work.

### Grant Support

This work was supported by Crohn’s and Colitis Foundation (CCF) Career Development Award (SZS), R01-ES024983 from NIEHS (SZS and TSF), 1R01DK104828-01A1 from NIDDK (SZS and TSF), P01-DK094779-01A1 from NIDDK (SZS), P30-DK034987 from NIDDK (SZS), 1-16-ACE-47 ADA Pathway Award (PS), UNC Nutrition Obesity Research Center Pilot & Feasibility Grant P30DK056350 (PS), CCF PRO-KIIDS NETWORK (SZS and PS), UNC CGIBD T32 Training Grant from NIDDK (JBB), T32 Training Grant (5T32GM007092-42) from NIGMS (MH), SHARE from the Helmsley Trust (SZS). The UNC Translational Pathology Laboratory is supported in part, by grants from the National Cancer Institute (3P30CA016086), the UNC University Cancer Research Fund (UCRF) (PS).

## References

1. Weiser M et al. Molecular classification of Crohn’s disease reveals two clinically relevant subtypes. Gut 2016;gutjnl-2016-312518.

2. Jostins L et al. Host-microbe interactions have shaped the genetic architecture of inflammatory bowel disease‥ Nature 2012;491(7422):119–24.

3. Cleynen I et al. Inherited determinants of Crohn’s disease and ulcerative colitis phenotypes: a genetic association study.. Lancet (London, England) 2016;387(10014):156–67.

4. Haberman Y et al. Pediatric Crohn disease patients exhibit specific ileal transcriptome and microbiome signature.. J. Clin. Invest. 2014;124(8):3617–33.

5. Kugathasan S et al. Prediction of complicated disease course for children newly diagnosed with Crohn’s disease: a multicentre inception cohort study. Lancet [published online ahead of print: March 2017]; doi:10.1016/S0140-6736(17)30317-3

6. McKenna LB et al. MicroRNAs control intestinal epithelial differentiation, architecture, and barrier function.. Gastroenterology 2010;139(5):1654–64, 1664.e1.

7. Peck BCE et al. Functional Transcriptomics in Diverse Intestinal Epithelial Cell Types Reveals Robust MicroRNA Sensitivity in Intestinal Stem Cells to Microbial Status.. J. Biol. Chem. 2017;292(7):2586–2600.

8. Chapman CG, Pekow J. The emerging role of miRNAs in inflammatory bowel disease: a review. Therap. Adv. Gastroenterol. 2015;8(1):4–22.

9. Pekow JR, Kwon JH. MicroRNAs in inflammatory bowel disease.. Inflamm. Bowel Dis. 2012;18(1):187–93.

10. Peck BCE et al. MicroRNAs Classify Different Disease Behavior Phenotypes of Crohn’s Disease and May Have Prognostic Utility.. Inflamm. Bowel Dis. 2015;21(9):2178–87.

11. Béres NJ et al. Altered mucosal expression of microRNAs in pediatric patients with inflammatory bowel disease. Dig. Liver Dis. [published online ahead of print: 2016];(2016). doi:10.1016/j.dld.2016.12.022

12. Lin J et al. Novel specific microRNA biomarkers in idiopathic inflammatory bowel disease unrelated to disease activity.. Mod. Pathol. 2014;27(4):602–8.

13. Kim D et al. General rules for functional microRNA targeting. Nat. Genet. [published online ahead of print: October 24, 2016]; doi:10.1038/ng.3694

14. Baran-Gale J, Fannin EE, Kurtz CL, Sethupathy P. Beta Cell 5’-Shifted isomiRs Are Candidate Regulatory Hubs in Type 2 Diabetes. PLoS One 2013;8(9). doi:10.1371/journal.pone.0073240

15. Uhlen M et al. Tissue-based map of the human proteome. Science (80-). 2015;347(6220):1260419–1260419.

16. Comelli EM et al. Biomarkers of human gastrointestinal tract regions. Mamm. Genome 2009;20(8):516–527.

17. Rutgeerts P et al. Predictability of the postoperative course of Crohn’s disease.. Gastroenterology 1990;99(4):956–63.

18. Zachos NC et al. Human Enteroids/Colonoids and Intestinal Organoids Functionally Recapitulate Normal Intestinal Physiology and Pathophysiology.. J. Biol. Chem. 2016;291(8):3759–66.

19. Perou CM et al. Molecular portraits of human breast tumours. Nature 2000;406(6797):747–752.

20. Tomczak K, Czerwińska P, Wiznerowicz M. The Cancer Genome Atlas (TCGA): an immeasurable source of knowledge.. Contemp. Oncol. (Poznan, Poland) 2015;19(1A):A68–77.

21. Lenz G et al. Molecular subtypes of diffuse large B-cell lymphoma arise by distinct genetic pathways.. Proc. Natl. Acad. Sci. U. S. A. 2008;105(36):13520–5.

22. Verhaak RGW et al. Integrated genomic analysis identifies clinically relevant subtypes of glioblastoma characterized by abnormalities in PDGFRA, IDH1, EGFR, and NF1.. Cancer Cell 2010;17(1):98–110.

23. Getz G et al. Integrated genomic characterization of endometrial carcinoma. Nature 2013;497(7447):67–73.

24. Collisson EA et al. Comprehensive molecular profiling of lung adenocarcinoma. Nature 2014;511(7511):543–550.

25. Hoadley KA et al. Tumor Evolution in Two Patients with Basal-like Breast Cancer: A Retrospective Genomics Study of Multiple Metastases. PLOS Med. 2016;13(12):e1002174.

26. Glocker E-O et al. Inflammatory bowel disease and mutations affecting the interleukin-10 receptor.. N. Engl. J. Med. 2009;361(21):2033–45.

27. Ostrowski J et al. Genetic architecture differences between pediatric and adult-onset inflammatory bowel diseases in the Polish population. Sci. Rep. 2016;6(1):39831.

28. Jakobsen C et al. Differences in phenotype and disease course in adult and paediatric inflammatory bowel disease - a population-based study. Aliment. Pharmacol. Ther. 2011;34(10):1217–1224.

29. Duricova D et al. Age-related differences in presentation and course of inflammatory bowel disease: an update on the population-based literature. J. Crohn’s Colitis 2014;8(11):1351–1361.

30. De Greef E et al. Diagnosing and treating pediatric Crohn’s disease patients: is there a difference between adult and pediatric gastroenterologist’s practices? Results of the BELCRO cohort.. Acta Gastroenterol. Belg. 2014;77(1):25–9.

31. Rosen MJ et al. Mucosal Expression of Type 2 and Type 17 Immune Response Genes Distinguishes Ulcerative Colitis From Colon-Only Crohn?s Disease in Treatment-Naive Pediatric Patients. Gastroenterology 2017;152(6):1345–1357.e7.

32. Cichon C, Sabharwal H, Rüter C, Schmidt MA. MicroRNAs regulate tight junction proteins and modulate epithelial/endothelial barrier functions. Tissue Barriers 2014;2(4):e944446.

33. Jovov B, Shaheen NJ, Orlando GS, Djukic Z, Orlando RC. Defective barrier function in neosquamous epithelium.. Am. J. Gastroenterol. 2013;108(3):386–91.

34. Sreedharan L et al. MicroRNA profile in neosquamous esophageal mucosa following ablation of Barrett’s esophagus.. World J. Gastroenterol. 2017;23(30):5508–5518.

35. Lin W-B et al. MicroRNA profiling of the intestine during hypothermic circulatory arrest in swine.. World J. Gastroenterol. 2015;21(7):2183–90.

36. Gao W et al. A systematic analysis of predicted MiR-31-targets identifies a diagnostic and prognostic signature for lung cancer. Biomed. Pharmacother. 2014;68(4):419–427.

37. Tian Y et al. Stress responsive miR-31 is a major modulator of mouse intestinal stem cells during regeneration and tumorigenesis. Elife 2017;1–30.

38. Dobin A et al. STAR: ultrafast universal RNA-seq aligner. Bioinformatics 2013;29(1):15–21.

39. Patro R, Duggal G, Love MI, Irizarry RA, Kingsford C. Salmon provides fast and bias-aware quantification of transcript expression. Nat. Methods 2017;14(4):417–419.

40. Love MI, Huber W, Anders S. Moderated estimation of fold change and dispersion for RNA-seq data with DESeq2.. Genome Biol. 2014;15(12):550.

41. Kanke M, Baran-Gale J, Villanueva J, Sethupathy P. miRquant 2.0: an Expanded Tool for Accurate Annotation and Quantification of MicroRNAs and their isomiRs from Small RNA-Sequencing Data doi:10.2390/biecoll-jib-2016-307

42. Kamada N et al. Unique CD14 intestinal macrophages contribute to the pathogenesis of Crohn disease via IL-23/IFN-gamma axis.. J. Clin. Invest. 2008;118(6):2269–80.

